# Predicting phenotypes from genetic, environment, management, and historical data using CNNs

**DOI:** 10.1101/2021.05.27.446033

**Authors:** Jacob D. Washburn, Emre Cimen, Guillaume Ramstein, Timothy Reeves, Patrick O’Briant, Greg McLean, Mark Cooper, Graeme Hammer, Edward S. Buckler

## Abstract

Predicting phenotypes from genetic (G), environmental (E), and management (M) conditions is a long-standing challenge with implications to agriculture, medicine, and conservation. Most methods reduce the factors in a dataset (feature engineering) in a subjective and potentially oversimplified manner. Convolutional Neural Networks (CNN) can overcome this by allowing the data itself to determine which factors are most important. CNN models were developed for predicting agronomic yield from a combination of replicated trials and historical yield survey data. The results were more accurate than standard methods when tested on heldout G, E, and M data (r=0.5 vs r=0.4), and performed slightly worse than standard methods when only G was held out (r=0.74 vs r=0.78). Pre-training on historical data increased accuracy by 1-36% compared to trial data alone. Saliency map analysis indicated the CNN has “learned” to prioritize many factors of known agricultural importance.

## Introduction

Prediction of phenotypes from a combination of environmental (E), genetic (G), and human-imposed (often referred to as management(M)) conditions has been a long standing challenge in biology and related fields (Messina et al. 2009, 2018; Technow et al. 2015; Cooper et al. 2016, 2021; Varshney et al. 2017; Washburn et al. 2020; Jarquin et al. 2021; Li et al. 2021). Most traditional approaches have focused on simplifying the problem by holding one or more factors constant (genetics, environment, management, or some subset within these) or accounting for some of the factors statistically as nuisance parameters. Methods have also generally been tailored to specific use cases and developed within a single siloed sub-discipline (Meuwissen et al. 2001; Jones et al. 2003; Holzworth et al. 2014, 2018; Crossa et al. 2017; Boote 2019). The methods used for prediction have also varied significantly, from fine-tuned mechanistic models designed to closely replicate physical and biological processes, to abstract statistical models with the goal of predicting phenotypes using the simplest mathematical function that results in the highest accuracy.

Mechanistic models such as crop growth models (CGM) rely on physiological equations validated through detailed experimentation and can make predictions with high accuracy when provided with high quality data (Holzworth et al. 2018; Soufizadeh et al. 2018; Hammer et al. 2020). However, the types of data and quality needed is prohibitively expensive and impractical for any more than a few cultivars (G) at a time. Geneticists and breeders have successfully used statistical models, primarily Best Linear Unbiased Prediction (BLUP) and genomic BLUP (GBLUP), for decades (Henderson 1975; Meuwissen et al. 2001; Gaffney et al. 2015). These methods have focused on prediction within the G space (sometimes called genomic prediction (GP)) but can be extended to incorporate information from E and/or M (Jarquín et al. 2014; Li et al. 2021). The downside to these methods is that they are not mechanistic in nature and require extensive feature engineering and data complexity reduction in order to avoid overfitting, limiting their interpretation, and perhaps accuracy. Conversely, GP methods are fairly robust to lower quality multi-environment trial data. Methods that incorporate both mechanistic and statistical components have recently been developed (Technow et al. 2015; Messina et al. 2018; Millet et al. 2019), and shown promise, but they still require data that is not easily collected by most research or breeding programs, and many aspects of the models have not been publicly released.

While current methods have had significant impacts on phenotype prediction and continue to yield important results in breeding and agronomy, they are each limited in the context of prediction across G, E, and M by one or more of the following: 1) a reliance on subjective feature engineering and factor reduction, 2) a need for expensive or low through-put data, and 3) the inability to utilize historical datasets not designed specifically for that method.

Machine learning approaches such as Convolutional Neural Networks (CNN) and others have the potential to overcome many of these current limitations. CNN’s for example were designed to utilize large parameter spaces and, in theory, estimate any continuous function regardless of its complexity, as long as enough examples are provided for training and the network is large enough (Zeng et al. 2016; Zhou 2020). This allows them to derive efficient representations of input data automatically, with limited feature engineering. CNNs are also extremely flexible to different input data types, and with large enough datasets (which is not always possible) and proper design they can perform robustly on low quality data (Yim and Sohn 2017; Qin et al. 2019). Additionally, while CNNs are often referred to as a “black box,” there are now well-demonstrated methods for dissecting and interpreting the processes within a given CNN model, and the features the model determines to be most important to prediction (Simonyan et al. 2013; Samek et al. 2019). Many of these interpretation tools, and even the machine learning methods themselves, are still in their infancy so it is likely they will continue to improve rapidly over the coming years.

Several authors have applied CNN and other machine learning models to plant phenotype prediction, but the quantity of data available in these studies was typically small when compared to the hundreds of thousands or even millions of data points used in more traditional CNN applications (e.g., image classification) (Montesinos-López et al. 2018, 2019; Pérez-Enciso and Zingaretti 2019; Zingaretti et al. 2020; Abdollahi-Arpanahi et al. 2020). Perhaps for this reason, phenotype prediction with CNNs has continued to rely on extensive feature engineering and factor reduction, and primarily used very shallow neural networks. A few attempts at using larger datasets with less feature engineering have been made, but these have not included a G component (Khaki et al. 2019; Shahhosseini et al. 2021).

In this study, CNN models for predicting grain yield were developed following a philosophy of maximum feature inclusion. Additionally, every attempt was made to include as many samples as possible in the dataset, to the extent of including historical samples with only partial data available, through transfer learning (using a pre-trained model as a starting point for model fitting). Extensive public maize field study data from The Genomes To Fields (G2F) Initiative, along with historical data from the United States Department of Agriculture - National Agricultural Statistics Service (USDA-NASS) and other governmental databases made using a very large dataset (>100,000 samples) possible (AlKhalifah et al. 2018; McFarland et al. 2020). On the other hand, all phenotypes, weather, soil, and management data came from publicly available sources and are the common types of data that a typical farmer, plant breeder, or agronomist could access or collect inexpensively today. While these models were developed in an agricultural system, the basic philosophy and results should be widely applicable across biology and other areas.

## Materials and methods

### Data mining and cleaning

Phenotypic, genetic, and field location and management data were obtained from The Genomes to Fields (G2F) initiative Genotype by Environment experiments from 2014 to 2017 (Gage et al. 2017; AlKhalifah et al. 2018; Falcon et al. 2020; McFarland et al. 2020). The phenotypic data were filtered to remove samples with yield values of zero, and to ensure that only samples with both genotypic and phenotypic data were included. Various other manipulations of the data were carried out in order to get them into a suitable format for the desired models (see bitbucket repository scripts). The genotype data for each cultivar were obtained directly from Anne Rogers based on the filtering process described in (Rogers et al. 2021). Principal components analysis was performed on the genotype data using TASSEL (Bradbury et al. 2007) and the first 30 principal components were used in the CNN. Methods to account for and visualize spatial field effects were also used to examine and filter the data, but these attempts appeared to have little impact on the results and so were not used in the final datasets and models.

Historical county level data were downloaded from the United States Department of Agriculture, National Agricultural Statistics Service for all the continental United States from 1970 to 2018. Weather data from each of these U.S. counties as well as from each of the G2F field sites were downloaded for the same time period from the DAYMET website (https://daymet.ornl.gov/) using custom scripts (see bitbucket repository) (Thornton et al. 2016).

The soil data used were pulled from the SSURGO Database as county-specific folders containing many correlated tables. Soil attributes available for any given county were highly variable so attributes with low representation were filtered from the set. Soil attributes were averaged across depths into three depth ranges.

### Training, validation, and test set development

Training, validation, and test sets were developed to examine four scenarios to represent those faced in agricultural prediction: the “strict G and E”, “practical G and E”, “G” and “E” holdout scenarios. For the strict G and E scenario (the scenario used for model structural optimization described below), all historical data from any year and any U.S. county within the G2F dataset were withheld from the training sets (Sup Fig S1). The G2F data were divided into 50 training, validation (for model structural and other optimizations), and test sets including the historical test set for cross-validation. Some field locations with less data were combined into a single fold, but most locations were left as separate folds. Each of the 49 G2F folds contained a training, validation, and test set. Cultivars in test sets were downsampled to ensure that each fold had the same number of cultivars represented, and the same number of total samples in the test set. Training sets were then formed from all locations that were not present in the test set and cultivars that were related to test set cultivars were excluded (based on k-means clusters on the genotypic data). In this testing scenario (strict G and E), different G2F years in the same location as a test set location were excluded from the training set. The validation set was derived from the training set by randomly sampling 5% of the training set and excluding those samples from the training set. Each of the other scenarios was designed in a similar manner except that sampling conditions about G and/or E were relaxed as detailed below.

For the practical G and E holdout scenario, the historical pre-training set was relaxed to include locations sampled in the G2F data during years previous to the G2F data. The same was true for the G2F data splitting, with a given training set being able to contain locations in the test set as long as they were not from the same year. Down sampling of cultivars to enforce strict testing set balance was also relaxed in this scenario to increase the number of samples in the sets.

The E holdout scenario was designed in a similar way to the two G and E holdout scenarios except that cultivars were not considered. Environments were defined similarly to the practical holdout set with the training sets being able to include the same field location as long as it was not in the same year. Down sampling and combining of some locations was also carried out in order to create more balanced test sets.

The G holdout scenario was similarly designed to the two G and E holdout scenarios except that environments were not held out. Down sampling of genotypes was again employed with the goal of creating balanced test sets. Because of the complications of doing this while also controlling for genetic relatedness of individuals, the final G holdout test sets were based on randomly chosen cultivars and the final training sets were constrained to exclude those cultivars.

All models were run in replicates, and the predicted test set values from the ten best performing replicates (based on the validation sets) was averaged to provide the final predicted value for each sample. Summary statistics were then calculated on a per fold basis and averaged to find the mean performance across all folds.

### CNN models

CNN models were created with 1D convolutional, dense, and dropout layers. Final models have three-streams, and these streams are concatenated to predict yield (Sup Fig S2). The first two streams (Weather and soil plus some field management available historically) were used for both historical and G2F data while the third stream (Genetics, fertility, and field management not available historically) was included only after pre-training. Since weather data were represented with a 1D time-series, 1D convolutional layers were used to extract weather-related features automatically in the weather stream. Because other streams were not a time series, they did not have convolutional layers and started with dense layers directly. We used a linear activation function in the last layer and Rectified Linear Unit (ReLU) in the rest. Mean squared error was used as the loss function.

CNN models have hyper-parameters that need to be selected and optimized for model accuracy. There are millions of possible hyper-parameter combinations, thus an efficient method is needed to select the best hyperparameter combination. We optimized the number of convolutional and dense layers, filter size, hidden layer size, number of kernels, pool size, strides, batch size, and Adam optimizer (used to train the model) parameters using the hyperopt package (Bergstra et al. 2013) based on validation set accuracy. The CNN models were built and run using the tensorflow.keras module version 2.1.0 in python 3.6.9 (Chollet 2015; Abadi et al. 2016b, a). The final model specifications can be found in the bitbucket repository.

### Saliency Maps

Saliency maps (Shrikumar et al. 2017; Zintgraf et al. 2017) were computed from the saved models from each CNN run using the tf-keras-vis module in python (https://github.com/keisen/tf-keras-vis). All reported scores are based on training sets and were averaged across replicates and splits to obtain the final values shown in the figures.

### BLUP models

Genotypic values of cultivars were predicted by best unbiased linear prediction (BLUP) methods, under the following polygenic model (Jarquín et al. 2014):

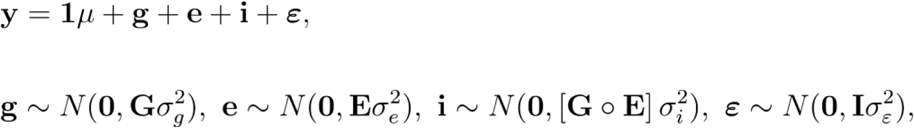

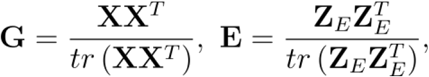

where **y** was the vector of cultivar phenotypes; **1***μ* is the grand mean of **y**. The vector **g** consisted of main genomic effects; **G** was the genomic relationship matrix, calculated from **X**, the (non-centered) matrix of allele counts at imputed SNPs; *tr* is the trace operator (sum of diagonals). The vector **e** consisted of main environmental effects; **E** was the environmental relationship matrix, calculated from **Z**_*E*_, the matrix of standardized environmental covariates (geographical location, management, soil properties, weather covariates), adjusted by their mean and scaled by their standard deviation. Weather covariates (day of year, day length, maximum and minimum temperatures, precipitation, radiation, vapor pressure, CTT) were included as averages over adjacent windows of width 3 (e.g., average temperature between day 1 and day 3). The vector **i** consisted of genome-by-environment interactions, as captured by **G**∘**E**, the Hadamard (elementwise) product of **G** and **E**. The vector **ε** consisted of independent and identically distributed errors. The BLUP model was fitted by restricted maximum likelihood (REML) using the R package qgg v1.0.4 (Rohde et al. 2020).

### Automated CGMs

Several approaches were developed and tested for calibrating APSIM version 7.10 crop growth models to new cultivars in an automated fashion based on limited phenotypic data (Holzworth et al. 2014, 2018). Calibration and testing of these approaches was carried out using the same training and test sets described above. The simplest automated method involved running APSIM on all available default maize cultivars within each of the training set environments. The best performing set of cultivar parameters for each actual cultivar in the dataset was then chosen based on the calculated error between observed and predicted yield values. In the scenarios in which G (cultivar) was held out, the genetic similarity matrix was used to assign the cultivar parameters of the most genetically similar training cultivar to the test cultivar. A second method was also used in which the default maize APSIM cultivars were again used as a starting point, but in this case, the parameter for growing degree days to flowering was adjusted to fit the known flowering dates of cultivars in the training set. Otherwise, the two methods were identical. A third method was also developed in which standard cultivar specific APSIM parameters were optimized to cultivars in the training set in an iterative fashion. This method was more computationally intensive and did not appear to work any better than the other two, so it was abandoned and not tested extensively.

## Results

All models were trained and tested under three basic holdout scenarios and some special cases. For most analyses E and M were held out jointly due to the limited amount of M data, and the extra computational time that would be required. However, the approach could be applied to separate E and M with little modification. In each scenario, a subset of G, E, and M, or combinations of the three, were withheld from the model and used as the test set (see Materials and Methods section for more details on training, validation, and test splits). Model performance was evaluated using a variety of metrics (Sup Table S1), but Pearson r values were used here to discuss the results, in keeping with the GP literature. In general, the trends seen in the Pearson r values were similar to those seen for other metrics such as root mean squared error (RMSE). Situations where this was not the case were noted. All results were based on test sets that included only data from the G2F experiments. The G2F data included over 38,000 data points and the historical data consisted of over 78,000 data points (Sup Table S2).

### The CNN model better extrapolates to unknown G, E, and M than other methods

Two scenarios were used to evaluate model performance while holding out G, E, and M. The scenarios used *k*-means clustering to hold out related cultivars. For the first scenario, which we named the “strict GEM holdout”, excluded all field locations in a given test set from the training set, including all data from that location in any other years. Although this scenario is not typical in agriculture, it does occur when new land is being used, or when records from the land do not exist. This scenario is also important for studying agriculture under climate change, as it tests the models’ ability to extrapolate to unknown environments and unknown cultivars, and represents a complete G, E, and M holdout. Under this scenario, all models performed poorly with Pearson r values ranging from 0.06 and 0.20 (Fig 1A). The GBLUP, CNN without historical data, GE BLUP and GxE BLUP methods were the poorest performers based on both Pearson r and RMSE (Sup Table 1). The pretrained CNN, which leveraged the historical survey data, performed the best (though still poorly), with an average Pearson r value of around 0.20. The RMSE of the pretrained CNN model represents a 15% decrease from that of the worst performing model, GBLUP (Table 1). The high standard deviation of all of the models in this scenario, makes it impossible to conclude with confidence that one model is better than another, but the CNN model with historical data performs the best on average.

**Figure 1.**
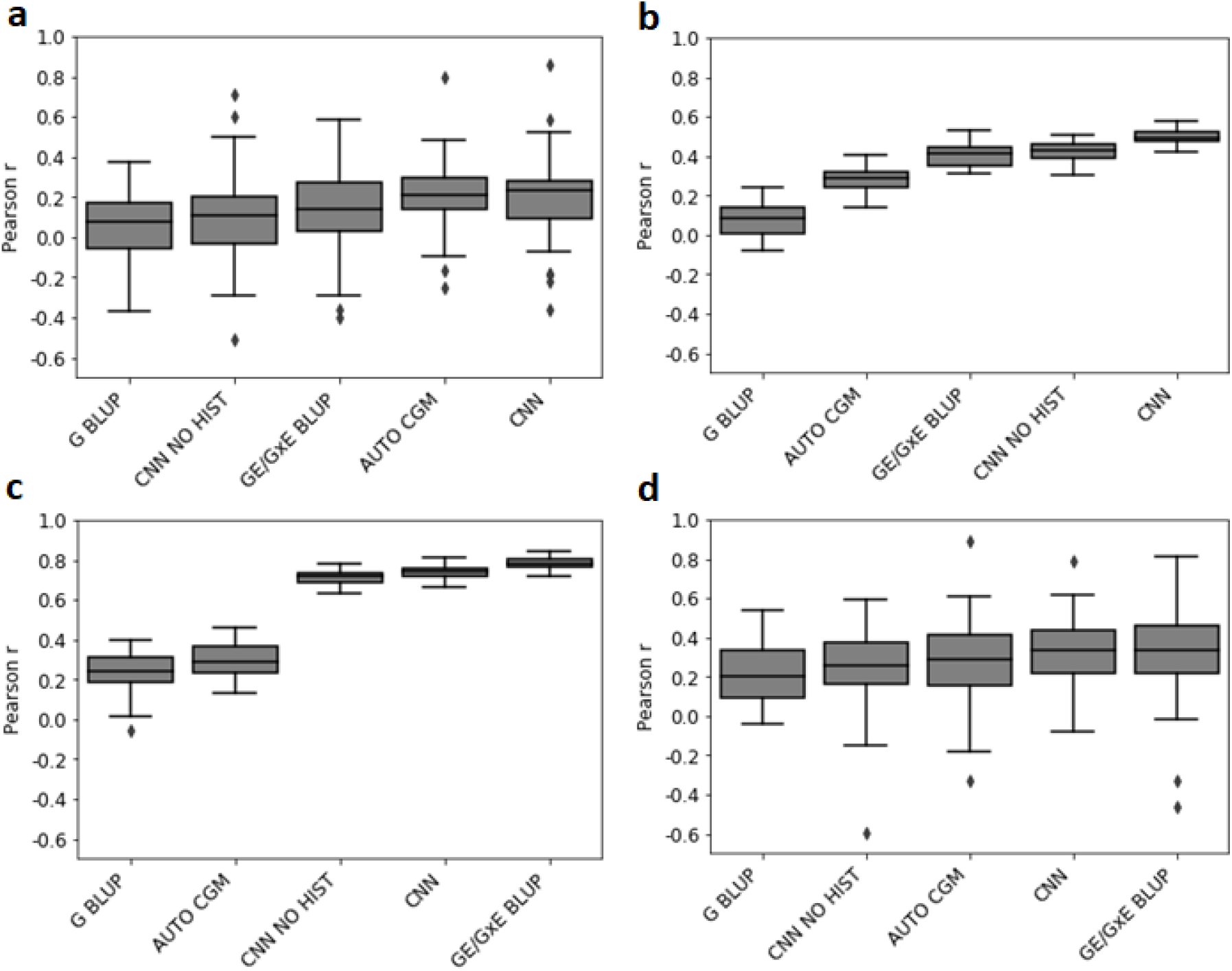
Pearson r values for different tested models in four scenarios. A) Strict genetic, environmental, and management hold out scenario with no previous year data given for held out environments and strictly balanced test sets. B) Practical genetic, environmental, and management hold out scenario with previous years data given. C) Genetic hold out scenario. D) Environmental and management hold out scenario. Box plot elements: center line, mean; box limits, upper and lower quartiles; whiskers, maximum and minimum with outliers excluded; points, outliers.

**Table 1.** Percentage increase or decrease in Pearson r and RMSE values between different methods and in different scenarios. For each scenario a comparison is shown between a Reference model (CNN without historical data, worst-performing model, best-performing BLUP model) and a Test model (CNN with historical data, best-performing model, best-performing CNN model). EM holdout is a training scenario in which a subset of both environment and management data are held out for testing. G holdout is a scenario where genetic data is held out for testing. The GEM scenarios hold out genetic, environmental, and management data in a “strict” or “practical” way, with environment and management data from other years in the same location being held out or not, respectively.

| Scenario | Comparison | Reference | Test | Pearson r % increase | RMSE % decrease |
| --- | --- | --- | --- | --- | --- |
| EM holdout | Without vs. With Historical | CNN NO HIST | CNN | 35.3 | 6.8 |
| EM holdout | Worst vs. Best | CGM | CNN | 9.5 | 26.6 |
| EM holdout | Best BLUP vs Best CNN | GE/GxE BLUP | CNN | N/A | 10.2 |
| G holdout | Without vs. With Historical | CNN NO HIST | CNN | 9.3 | 8.9 |
| G holdout | Worst vs. Best | CGM | GE/GxE BLUP | 166.7 | 47.5 |
| G holdout | Best BLUP vs Best CNN | GE/GxE BLUP | CNN | N/A | N/A |
| GEM practical | Without vs. With Historical | CNN NO HIST | CNN | 6.4 | N/A |
| GEM practical | Worst vs. Best | CGM | CNN NO G | 79.6 | 26.5 |
| GEM practical | Best BLUP vs Best CNN | GE/GxE BLUP | CNN NO HIST | 15.3 | 8.1 |
| GEM strict | Without vs. With Historical | CNN NO HIST | CNN | 1.1 | 2.0 |
| GEM stric | Worst vs. Best | CGM | E BLUP & CNN | 2.5 | 15.5 |
| GEM stric | Best BLUP vs Best CNN | GE/GxE BLUP | CNN | 53.2 | 3.9 |

The second G, E, and M) holdout scenario is here called the “practical GEM holdout” as it is meant to replicate the types of situations farmers, breeders, and researchers most commonly encounter. In this case, the training set was constructed in a similar way to the strict GEM scenario, except it included locations that were in the test set as long as the location data came from a different year than that being tested. The models performed with Pearson r values ranging from 0.08 to 0.50 (Fig 1B). The GBLUP, AUTO CGM, GE BLUP, and GxE BLUP models performed most poorly while the CNN with historical data performed the best (r ~ 0.50). The CNN with and without historical data performed more similarly based on RMSE values (Sup Table 1). The RMSE of the best performing model represents a 25% decrease from that of the worst performing.

When the models were trained under a “G holdout scenario” (training on all environmental and management data but with a portion of the cultivars withheld as a testing set) they performed with average Pearson r values across folds ranging from 0.24 to 0.78 (Fig 1C). The standard GBLUP and AUTO CGM models performed far worse than the other models in the G holdout scenario. When trained only on the G2F data set and corresponding environmental variables, the CNN model performed with an average r value of 0.68 but was boosted to a value of 0.74 when the model was pre-trained on historical data. The best performing model in the G holdout scenario was the GxE BLUP model (Jarquín et al. 2014) with an r value of 0.78. The RMSE of the best performing model represents a 47% decrease from that of the worst performing (Table 1).

Model performance under the “EM holdout scenario” (trained using all cultivars but withholding environmental and management data) resulted in average Pearson r values between 0.21 and 0.33 (Fig 1 B). The worst performers in this case were the G BLUP, CNN without historical data, and AUTO CGM. The CNN model with historical data training, and the GE BLUP model both performed relatively similarly on average, but all models had high standard deviations making it difficult to conclude which performed the best. The RMSE values gave a slightly different ordering for the performance of these methods, with the CNN with historical data method outperforming all others by a small margin (Sup Table 1). The RMSE of the best performing model represents a 27% decrease from that of the worst performing.

### Pre-training with historical data sometimes increased model accuracy

Historical maize yield and weather data by U.S. county are available going back decades, as well as low resolution soil and weather data across most of the continental U.S. (see Materials and Methods). Pre-training the CNN model based on these data and then fine-tuning the model using the modern G2F dataset (see Materials and Methods) resulted in increases in accuracy for some of the scenarios but little change for others (Table 1). For the G holdout scenario and the EM holdout scenario, historical pre-training resulted in 9% and 7% reductions in RMSE, respectively, in comparison to the same scenario without pre-training. For the two G and E holdout scenarios changes in RMSE were minimal. The CNN model was also run using the historical data as the only training, but still testing on the G2F data as above. In this case the model performed much worse than it did when trained on both the historical and G2F data, or on the G2F data alone (Sup Table 1). Together, these results indicate that pre-training with historical data improved prediction accuracy but was not itself sufficient for predicting yield in G2F trial data.

### What factors are most important in the CNN model?

Saliency maps were used to investigate the importance of each input factor in the model. In general, the holdout scenario had little impact on the relative importance scores for each model factor, but the use of historical pre-training had a major impact on the scores (Sup Fig S3). Precipitation, vapor pressure, and planting density were among the most important factors in the model independent of whether or not historical pre-training was used (Sup Fig S4). When summarized into overall categories (weather, soil, field, fertility, and genetics), the most important factors with historical pre-training were soil (35% of total importance score), weather (22%), genetics (20%), field (15%), and fertility (8%) (Fig 2a). Conversely, when historical data was not included in the model the genetic data was the most important (40%) followed by weather (23%), fertility (19%), soil (14%), and field data (3%) (Fig 2b). These differences are likely a direct result of not having genetics and fertility data in the historical set.

**Figure 2.**
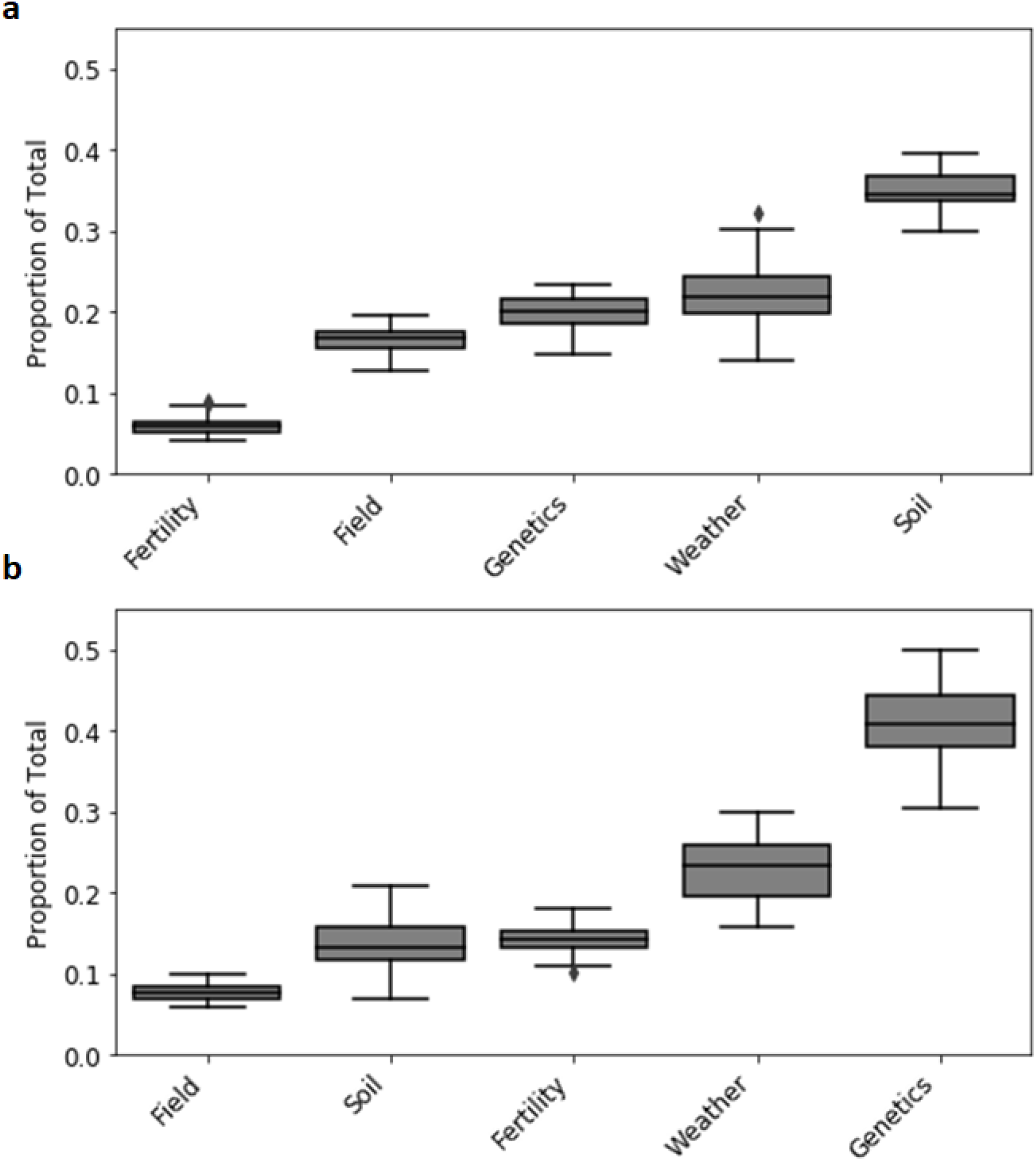
Saliency map score summaries displayed as a proportion of the total of all scores for each of the five main data type categories included in the models. A) With the use of historical data in model training. B) Without the use of historical data in model training. Box plot elements: center line, mean; box limits, upper and lower quartiles; whiskers, maximum and minimum with outliers excluded; points, outliers. All values based on the practical GEM scenario.

Weather-related factors were examined in more detail both across and within the growing season. The importance of weather-related factors did not change substantially across different training set scenarios, but it did change for some factors depending on the inclusion of historical data (Sup Fig S5). The most important weather-related factor in the model was precipitation, accounting for an average of 37%-42% of the total saliency map weather score in the practical G and E holdout scenario (Fig 3). The ranking of other weather factors changed somewhat depending on the use of historical pre-training (Fig 3). Combining maximum and minimum temperature together into a single factor resulted in temperature being the second most important factor regardless of historical pre-training (20%-26% on average). Vapor pressure is also one of the most important factors accounting for 11-18% of the score. Overall, factors related to humidity and temperature captured most of the signal about environments, as opposed to radiation and daylength.

**Figure 3.**
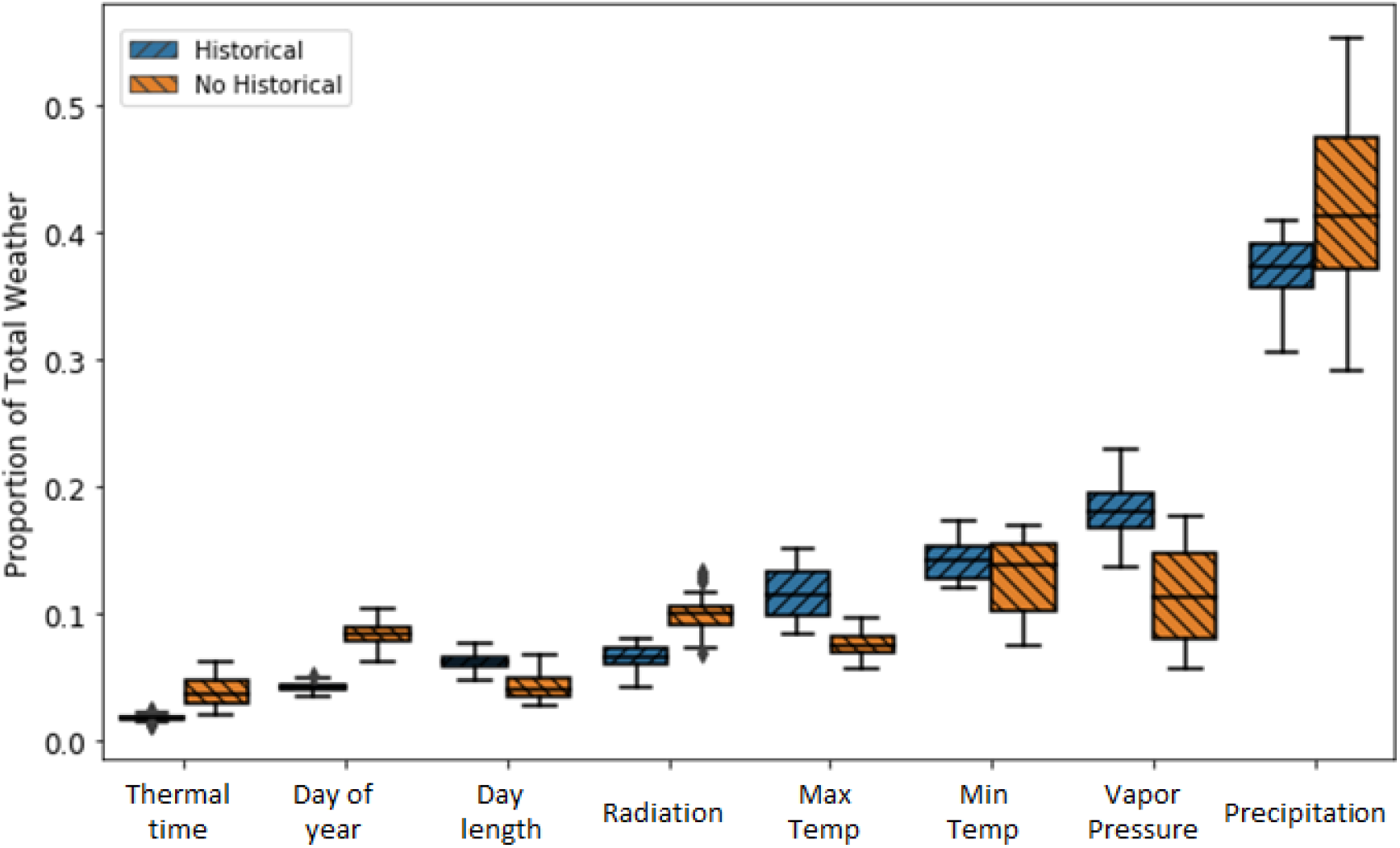
Saliency map score summaries displayed as a proportion of the total of all weather-related scores for each of the eight weather categories used in the models. Box plot elements: center line, mean; box limits, upper and lower quartiles; whiskers, maximum and minimum with outliers excluded; points, outliers. All values based on the practical GEM scenario.

The weather-related saliency scores on a daily basis are reasonably similar across all training scenarios except for the G and E holdout scenarios where historical pre-trained CNN models show some days in which the vapor pressure exceeds the saliency score for precipitation (Sup Figs S6-S7). As a general trend, precipitation had higher scores than other factors across the entire season regardless of pre-training (Fig 4). However, the importance of other weather-related factors increased with pre-training. Precipitation typically had its highest daily scores soon after planting, peaking around 5-15 post planting, and then having two smaller and flatter peaks around 50 and 110 days after planting. The other factors had similar trends to precipitation with the exception of vapor pressure and minimum temperature, both of which had their second peak shifted earlier at around 30-35 days post planting when pre-trained on historical data (Fig 4A). In the case of both the G alone and E alone holdout sets with historical pre-training this shift was also seen for maximum temperature (Sup Fig S6-S7). Therefore, pre-training affected the relative importance of weather-related factors, as well as time periods during the growing season.

**Figure 4.**
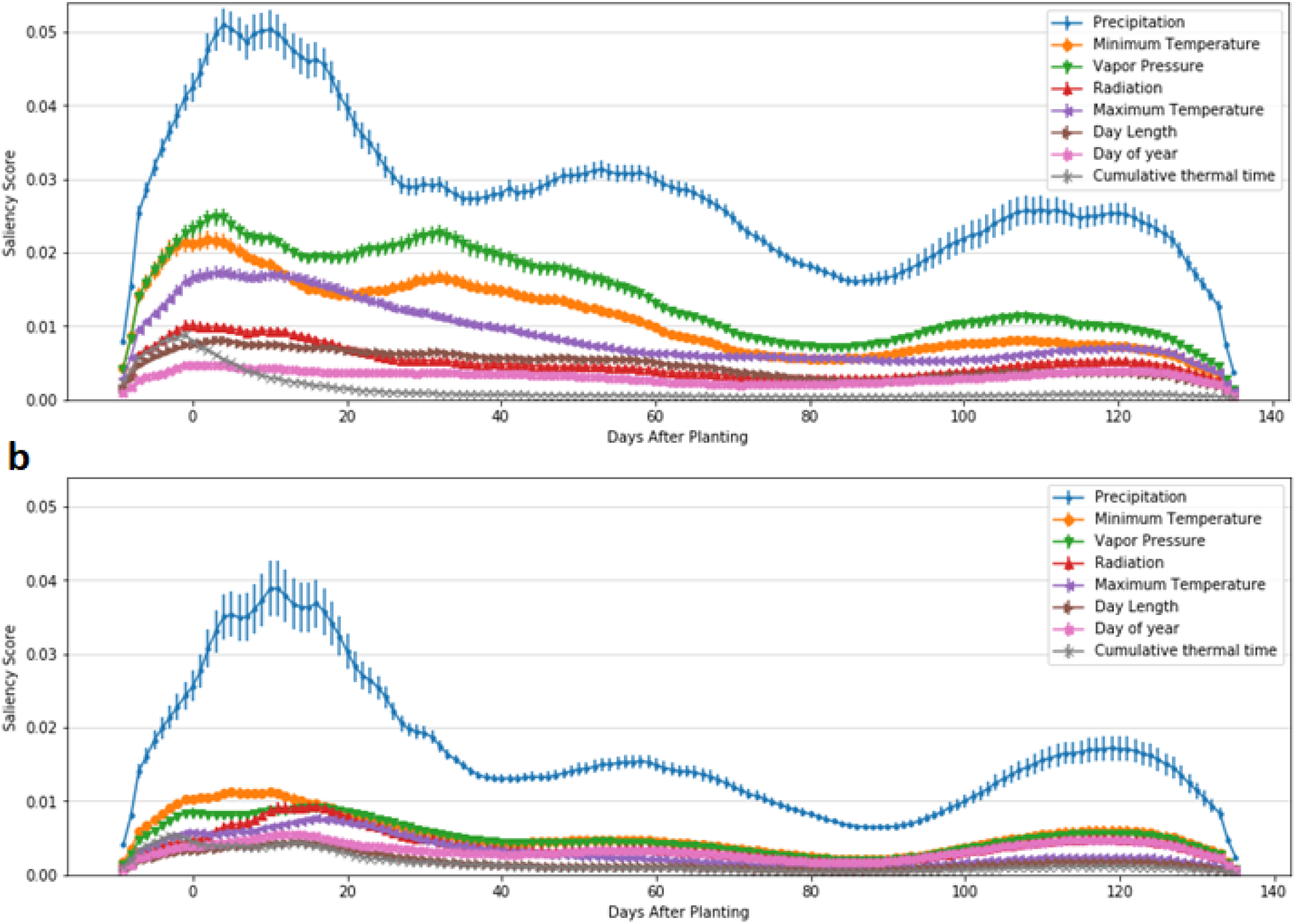
Daily Saliency map weather scores. Values are the average across all splits from the practical G and E holdout set with (A) and without (B) historical pre-training. Error bars are the standard error across splits.

Soil and fertility related factors with highest importance scores included soil electrical conductivity, calcium carbonate, saturated hydraulic conductivity (Ksat), gypsum content, and others (Sup Fig S8). Soil fertility and field management factors with the highest importance scores included planting density, irrigation, and percent sodium (Na) (Sup Figs S9-S10). Standard fertilizer components such as nitrogen (N), phosphorus (P), and potassium (K) also had substantial contributions when trained without historical data. In general, soil and management factors related to water use and plant density show the highest importance values, followed by NPK, and micronutrients.

To further dissect the importance of different factors, the models were run while excluding categories of factors (Sup Fig S11). Due to computational constraints, not all combinations of factors were tested, and updated models were not re-optimized. The exclusion of any single category (Soil, Fertility, Field/Management, Weather, or Genetics) from the model, under any of the training scenarios, generally resulted in a decrease in accuracy, but these decreases were typically small, and the removal of historical pre-training from the model usually resulted in larger reductions than the exclusion of any single category of factors (Sup Fig S11). However, training the model with only one factor included at a time generally resulted in poor model accuracies. Therefore, there is complementarity among categories of factors, even though their contribution to accuracy is not entirely distinct.

## Discussion

### Assumptions, sources of error, and complexity of the data and models

The results presented above should be considered in the context of the datasets used for analysis. The final Genomes to Fields (G2F) dataset here used was a combination of phenotype, environmental, and management data taken by hundreds of different individuals across more than 90 distinct location-by-year environments spanning much of the continental U.S. (see Materials and Methods). Each site and year was different in terms of the environmental, social, and technical conditions that influenced the planting, agronomic practices, data collection, and many other variables. Extensive measures were taken to reduce these errors and to account for them statistically, but there is no doubt that many factors remain unaccounted for even in the most complex models here developed. Because of a desire to include as many data points as possible in the models, very little data were removed through filtering.

It is also important to consider several decisions made during the construction of the CNN model that are likely to impact the results in different scenarios. The structure of the CNN model was optimized using validation data within the most challenging “strict G and E” holdout scenario (see Materials and Methods). The model structure was not optimized for any of the other scenarios, but simply trained on data from each of those scenarios. For that reason, the results presented above are likely underestimates of how well a CNN model could perform in those scenarios if the structure were optimized for them. Considerable effort was spent to develop training and test sets that would not bias the model or allow it to predict well based on extraneous factors. The strict G and E holdout scenario in particular was designed as a stringent scenario to minimize opportunities for a model to “memorize” details of the training set and still do well on the test set. While we cannot completely rule out the possibility that the model is gaining accuracy from extraneous factors, the saliency maps, and results from excluding different factors indicate this is unlikely.

### Value and trade-offs of using sparse historical data

The use of sparse, low-resolution historical data from commercial corn production over the past several decades, as well as additional low-resolution soil and weather data from national repositories, generally resulted in model accuracy increases (Table 1). The increases in r values were most pronounced in the E holdout scenario (35%) followed by the G holdout (9%), practical G and E (6%), and hard G and E (1%) scenarios. It is interesting that the historical data were most helpful in the E holdout and G holdout scenarios. The explanation for this may be at least partially attributed to the fact that these scenarios have more training data to begin with (they only hold out G or E). The historical data were also held out to a lower extent in these scenarios, so the models were trained on much larger datasets than they could be without historical data. Historical data pre-training also appeared anecdotally to increase model stability across training epochs.

The saliency map results indicate that models trained without the historical data make less use of weather and soil data than those trained with historical data. One might reasonably conclude that the approximately 90 examples of weather and soil data corresponding to the G2F trials are simply not enough examples for the model to learn how these factors are correlated with yield. This potential lack of information about environments might be all the more limiting since empirical models like CNNs rely completely on the data they are given, as opposed to mechanistic models which use known relationships between factors and plant growth. In that case, an argument can be made that a model pre-trained on thousands of historical examples of the relationship between weather, soil, and yield is likely to be more transferable to new datasets. On the other hand, the lack of genetic, field/management, and high-resolution soil and fertility data in the historical set introduces a potential bias to the model because it has more opportunities to learn about weather and soil features, and therefore may overstate their importance. With the current dataset, we cannot test this hypothesis since we do not know how the model would perform, and what factors would be most important, in the hypothetical case where we had high-resolution genetic, fertility, and other data for each historical data point.

### Importance of different factors

Arguments surrounding the relative importance of genetics, environment, and management have occurred for many decades. One would be hard pressed to conclude from the literature, or the results of this project, that any one factor is the most important or should be studied at the exclusion of the others. Additionally, management factors were largely underrepresented in the G2F dataset. That said, regardless of the training scenario, several factors consistently ranked highest in their importance scores. One clear conclusion, which is supported by both the literature and the current state of knowledge in maize production, is that planting density is a critical factor in determining maize yield (Duvick 2005). Precipitation also showed an outsized importance score when compared to most other factors in the model. The importance of precipitation in plant agriculture is well-known and obvious, and it has long been regarded, along with soil properties that change water availability to the plant, as one of the most critical factors in crop growth models (Barnett and Thompson 1982; Riha et al. 1996; Togliatti 2017). Our findings further validate and reinforce the idea that both soil properties and weather parameters related to water are of critical importance in predicting and/or understanding plant growth across environments. Fertilizer-related values (particularly NPK) were not as important as we had expected, but this is likely due to the fact these values do not exist in the historical data, and even the G2F experiments were mostly performed with relatively low amounts of fertilizer compared to commercial production.

Weather factors associated with temperature have also long been considered of significant importance in predicting plant growth and development (Togliatti 2017; Tollenaar et al. 2017). In this study of the G2F data set, daily minimum and maximum temperature were both of high importance to the CNN model. Cumulative thermal time, calculated from the maximum and minimum temperature, is considered a critical phenological component in crop modeling, and was calculated and included as a factor in the models trained here. Thermal time had the lowest importance score of any weather factor in the CNN model. This is likely due to the strong relationship between temperature and thermal time, and the ability of CNNs to produce their own functions of temperature (and other raw input data) for more accurate predictions. It appears the CNN model was able to gain more value from the two temperature factors alone than from their reduced representation as thermal time. Atmospheric vapor pressure was also an important factor (11-18%). Vapor pressure is related to effects of both water availability and temperature, and is known to play an important role in the plants’ ability to transpire, draw in nutrients, and carry out photosynthesis (Rawson et al. 1977; Lobell et al. 2013; Yuan et al. 2019). Radiation, day length, and day of year all had relatively low importance scores, but not low enough to be assumed insignificant. These factors also generally have close relationships with temperature, which may have reduced their utility in the model.

Some of the most important soil factors included soil electrical conductivity (EC), gypsum content, calcium carbonate, saturated hydraulic conductivity (Ksat), and cation exchange capacity. EC is a proxy measurement associated with water-soluble salt concentrations in the soil. The concentration of salts in the soil can make it more or less difficult for the plant to absorb water because it changes the osmotic pressure. Gypsum content in the soil has been shown to be important to plant growth in many ways: it is a source of both calcium and sulfur, it aids in soil water absorption, it can improve root length density, it can improve the physical structure of soils, and the benefits of gypsum amendments can be very long lasting (Shamshuddin et al. 1991; Wallace 1994; Toma et al. 1999; Chaganti et al. 2019; Macana et al. 2020). Gypsum had a high percentage of missing data across the locations. Moreover, its missingness may not be random (and may carry some information). It is possible that the model inflated gypsum’s importance for predictions, particularly in the pretrained CNN models. This possibility was examined by running models without Gypsum, other soil parameters with high missing data, and without soil at all (Sup Fig S8). Removal of these factors did reduce model accuracies in some cases, but not in such a substantial way as to suggest that the CNN model was overfit based on the presence or absence of data for these factors. Still, a conclusion that gypsum is more important than other soil factors in the model is probably not warranted based on this and previous studies. Ksat is a physical property related to how quickly water is transmitted through soil pores and has long been recognized as an important factor in soil-plant water relations and crop growth modeling (Kunze et al. 1968; Castellini et al. 2021). Cation exchange capacity is a measure of the soil’s ability to hold cations. It is related to the soil’s ability to hold fertilizers as opposed to leaching them into the groundwater, and therefore has potential impacts on plant growth, sustainability, and pollution (Kaiser et al. 2008). It is important to note that these factors were obtained from the SSURGO database and are therefore static factors that are the same across all years for a single location. In reality, many of these factors have the potential to vary substantially over time at any single field location. Because the factors were input as static across years, they also have the potential to artificially inflate our results in model scenarios where all years at the same location are held out. We do not see evidence of this occurring in our results, but it is a potential drawback of using static soil parameters as was done here.

While CGMs provide the means to integrate the effects on yield of these static variables in a more dynamic and biologically meaningful manner, the sensitivity of responses to water limitation likely requires quality in soil and crop data beyond that available in the G2F and historical data sets, in order to support robust CGM prediction. Messina et al., (Messina et al. 2019) showed the extreme sensitivity of maize yield to small differences in water stress level at flowering. Cooper et al., (Cooper et al. 2016) demonstrated the need to incorporate variation for soil depth (water-holding capacity) within experimental fields as an uncertainty factor in fitting procedures to capture the added value possible in fitting GP models using advanced Bayesian computation methods to link CGM-GP. The trade-off between data quantity and data quality remains an issue of interest as limited additional effort to capture meaningful data (e.g. soil water holding capacity estimates) may enable much more effective use of CGM, and also CNN. Advanced methodologies provide a path for integrating physiological understanding of variation in elite germplasm into a breeding program without the cost and complexity associated with measuring all of the relevant traits on all of the entries within all of the environments sampled (Cooper et al. 2016; Messina et al. 2018, 2019).

Genetic factors also played a strong role in prediction accuracy. However, because these models were not designed with the genetic resolution necessary to pinpoint specific genetic factors, conclusions on which genetic factors were most important are not straightforward. Here, genotypic data was incorporated into the CNN as principal components, to capture broad patterns in genetic variability. Further developments are needed to use complex neural network architectures for genetic inference (Demetci et al. 2020; Zhao et al. 2021).

### Conclusions and future directions

The CNN model approach with daily weather inputs and historical data was found to be capable of predicting maize yield at similar or higher accuracies than the standard BLUP approaches commonly used in genetics and breeding. This indicates the model is able to gain predictive value from the additional features and complexity it can take as input and suggests further study and development of CNN models in phenotype prediction is warranted. Automated mechanics models, like the CGMs here developed and tested, appear unable to perform well given the low overall quality, and the types of data used in this study. Higher quality data, more data of the specific types commonly used in CGMs, and/or better automation/integration methods will be needed in order to use CGM’s effectively across many cultivars in the scenarios presented here. Additional and low-cost attention to improve data quality may well favor use of CGMs in the long run. Historical survey data, despite being of low-resolution and quality, contains valuable information that can be utilized by models to improve accuracy through pre-training and transfer learning. Data types that are typically underrepresented in field experiments (such as soil data, and weather across time) may be key to developing models that more realistically represent the importance of these factors.

Although CNN models are often described as being difficult to interpret, the results here demonstrate that our model reconstructed the importance of various agronomic factors in a way that is consistent with the literature. This indicates that given sufficient data of the right kinds, CNN and other deep learning models may have some utility in the development of mechanistic hypotheses for further testing and validation. Despite the available tools for interpretation (and the many new tools that will likely come), deep learning methods are fundamentally based on empirical data, and will likely never reach the level of interpretation possible in CGMs. Additionally, deep learning models do not provide some types of interpretive data (for example heritability, additivity, and dominance scores) commonly generated by standard statistical approaches. Moreover, their assumptions, results, and potential pitfalls deserve greater attention, before they can be widely adopted by the community.

Future development and testing of machine and deep learning approaches in phenotype prediction would benefit from larger, more balanced agricultural datasets, and/or methods for better validating these models using the datasets currently available. The questions of how best to incorporate data into the models, what model architectures to use, and what data types are most important need significantly more exploration. Additionally, the possibility of incorporating known physiological, chemical, or other mechanisms into machine and deep learning models should be explored.

## Declarations

### Availability of data and material (data transparency)

All data is publicly available at: https://www.genomes2fields.org/resources/, https://daymet.ornl.gov/, or https://websoilsurvey.sc.egov.usda.gov/App/WebSoilSurvey.aspx

### Code availability

All code is available at (BitBucket repository link)

### Authors’ contributions

JW, EC, GR, GH, MC, and EB conceived the project. JW, EC, and GR implemented the CNN and BLUP methods. JW, TR, and RO developed the data mining tools. JW and GM developed and performed the APISM methods. All authors contributed to the writing and revision of the manuscript text.

### Key Message

Convolutional Neural Networks (CNNs) can perform similarly or better than standard genomic prediction methods when sufficient genetic, environmental, and management data is provided.

## Acknowledgments

This material is based upon work supported by the United States National Science Foundation (NSF) NSF Postdoctoral Research Fellowship in Biology under Grant No. 1710618 (JDW). Additional support comes from the United States Department of Agriculture - Agricultural Research Service. Anna Rogers and James Holland provided pre-filtered genotype and phenotype data for the Genomes to Fields Initiative.

