## Supplemental Figures for "Predicting phenotypes from genetic, environment, management, and historical data using CNNs"

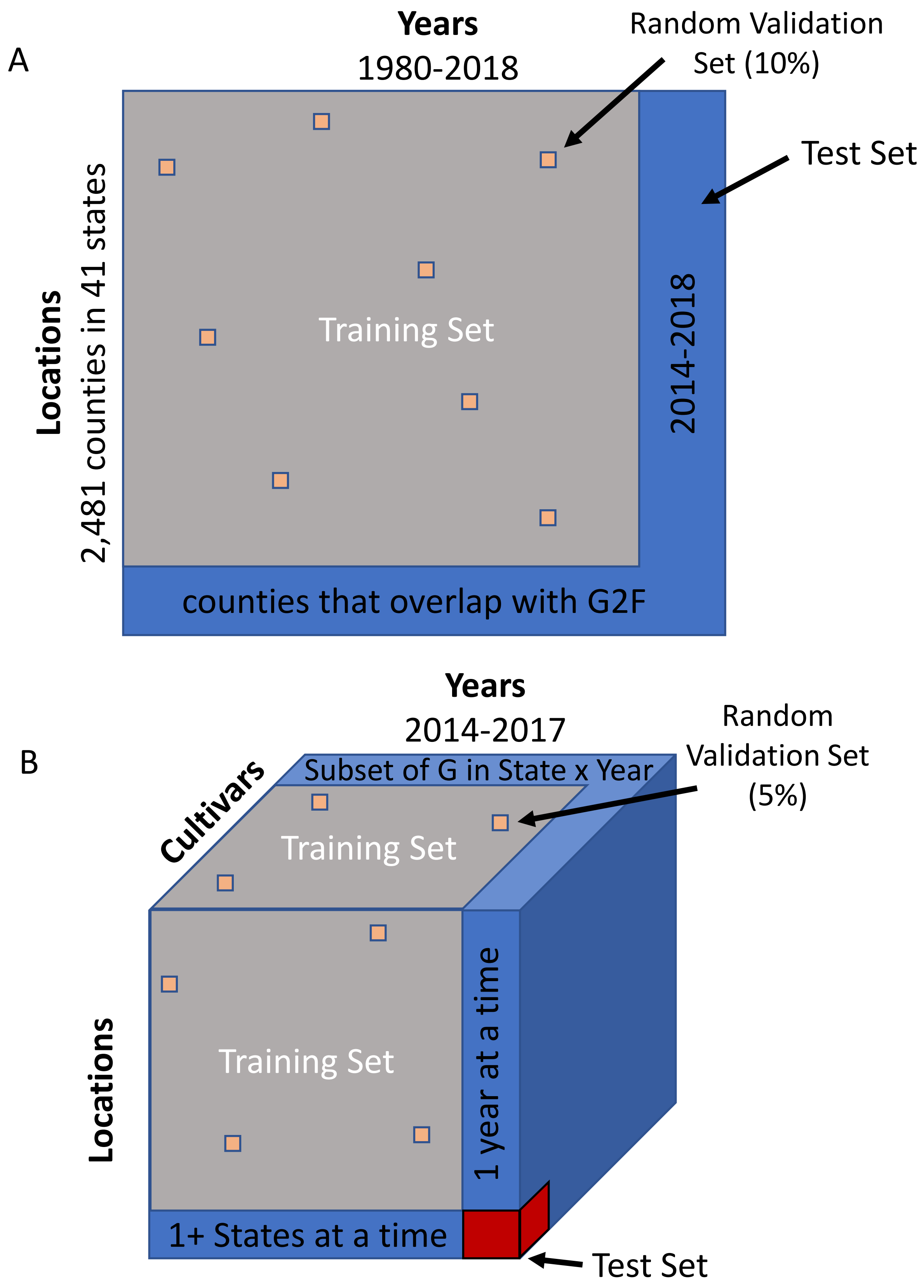


Supplemental Figure S1. Illustrations of how training, validation, and testing sets were determined for A) The historical dataset, and B) the Genomes to Fields (G2F) data set. Illustrations are based on the GEM strict scenario.

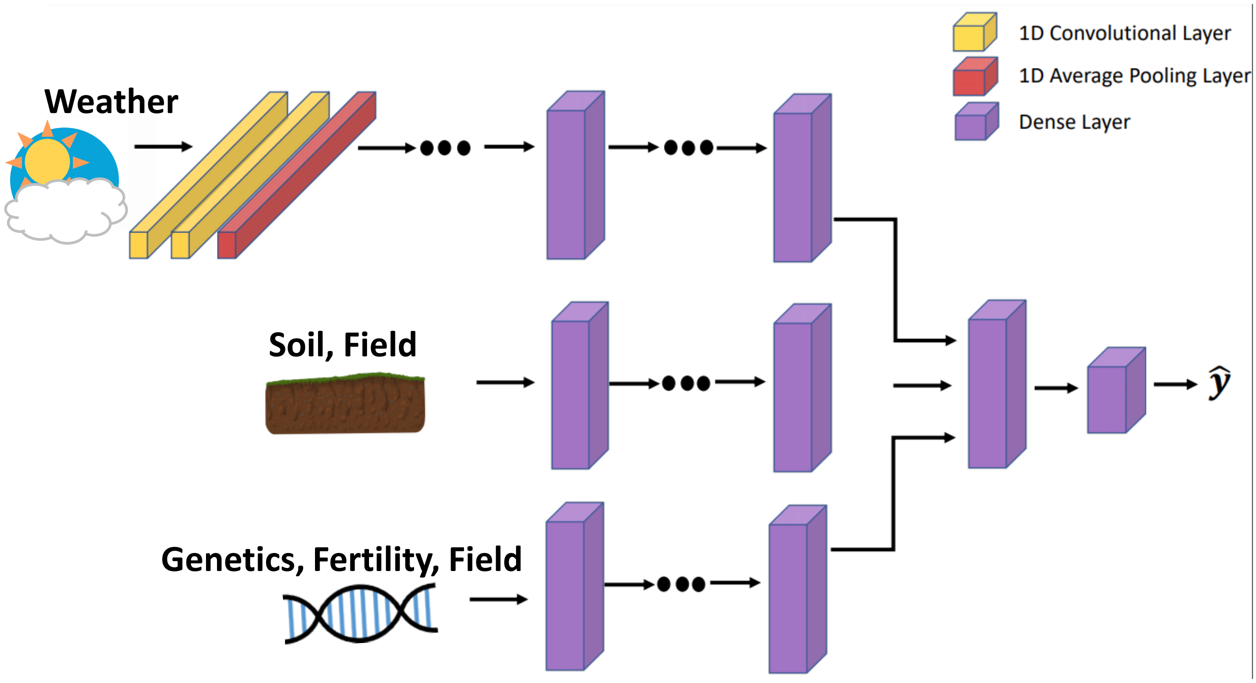


Supplemental Figure S2. Illustration of the basic structure of the neural network used for predicting yield from genetic, environmental, and management data. Some field management values were included with soil (if available in historical data), and some were included with Genetics and Fertility (if not available in historical data).

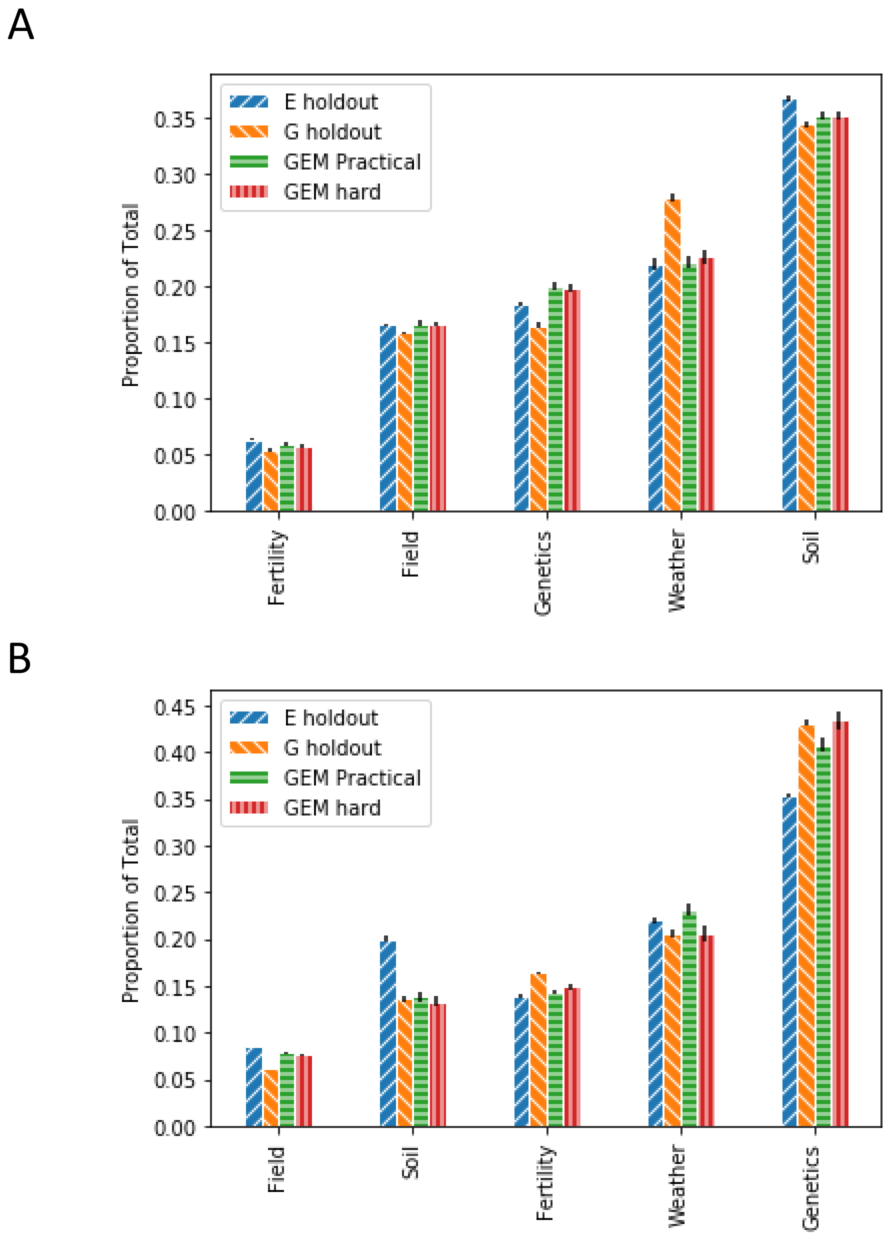


Supplemental Figure S3. Proportion of total saliency map scores attributable to different categories of factors. A) After pre-training on historical data, B) No pre-training on historical data.

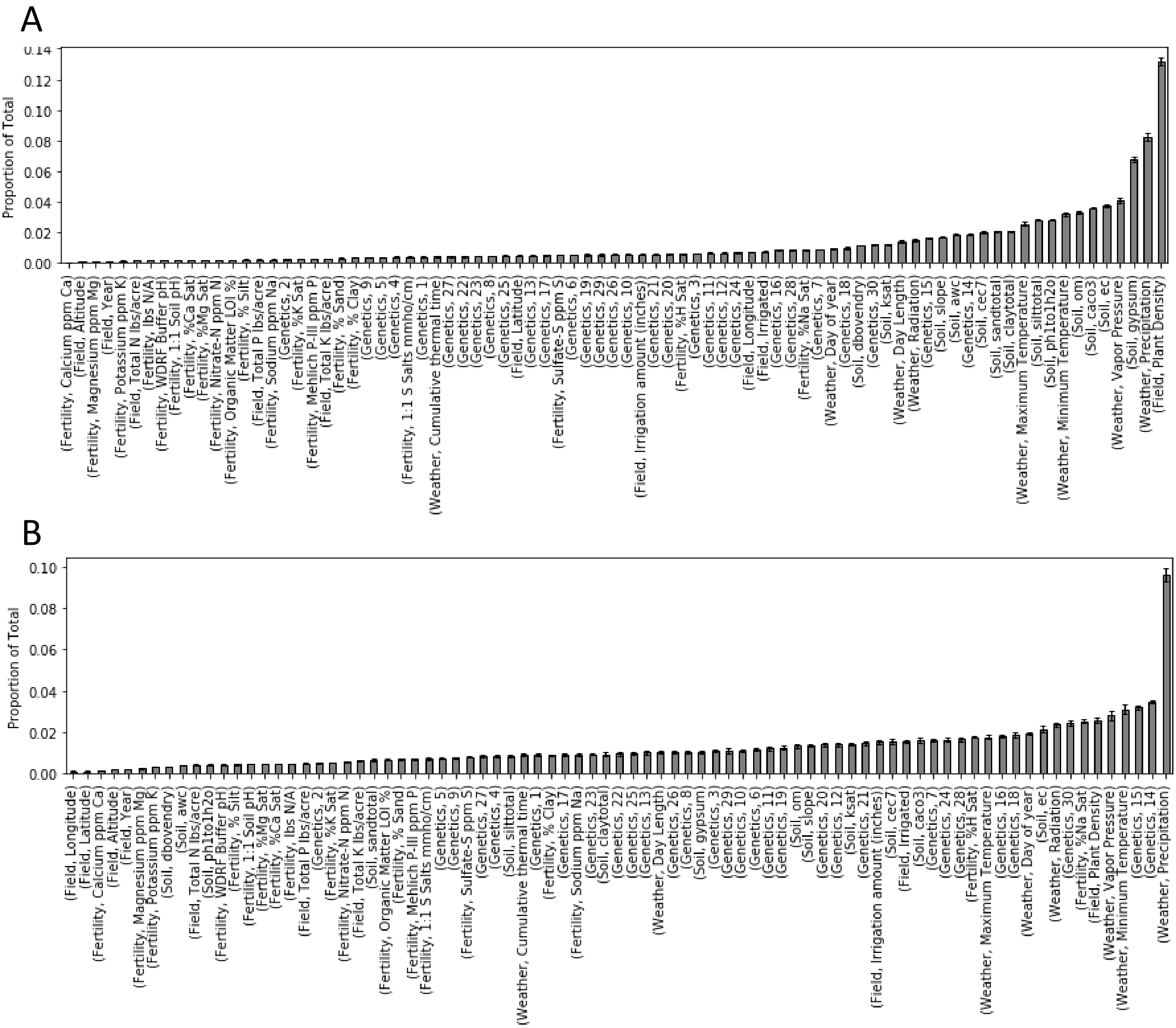


Supplemental Figure S4. Proportion of total saliency map scores attributable to different model factors. A) After pre-training on historical data, B) No pre-training on historical data.

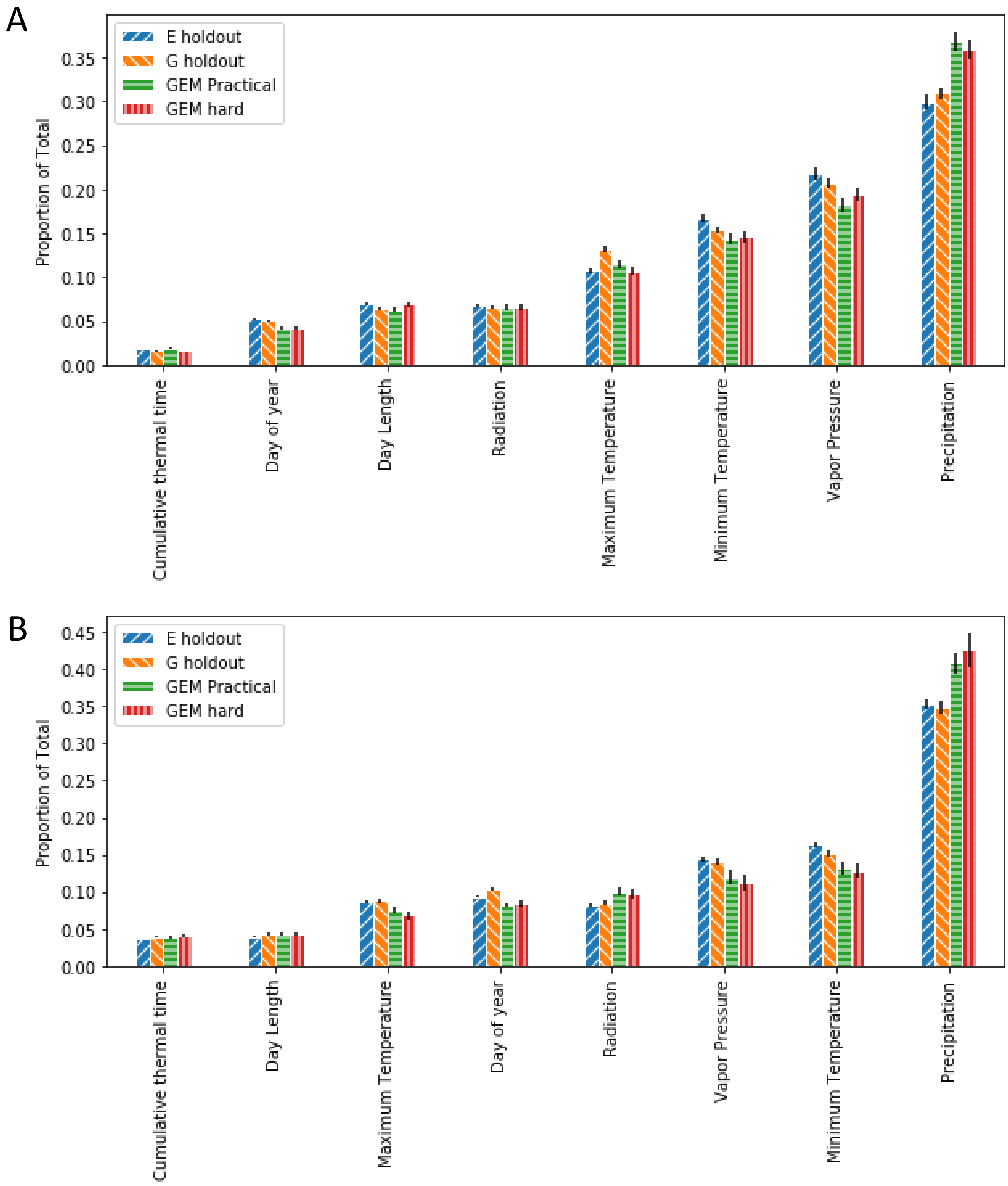


Supplemental Figure S5. Proportion of weather-related saliency map scores attributable to different weather factors. A) After pre-training on historical data, B) No pre-training on historical data.

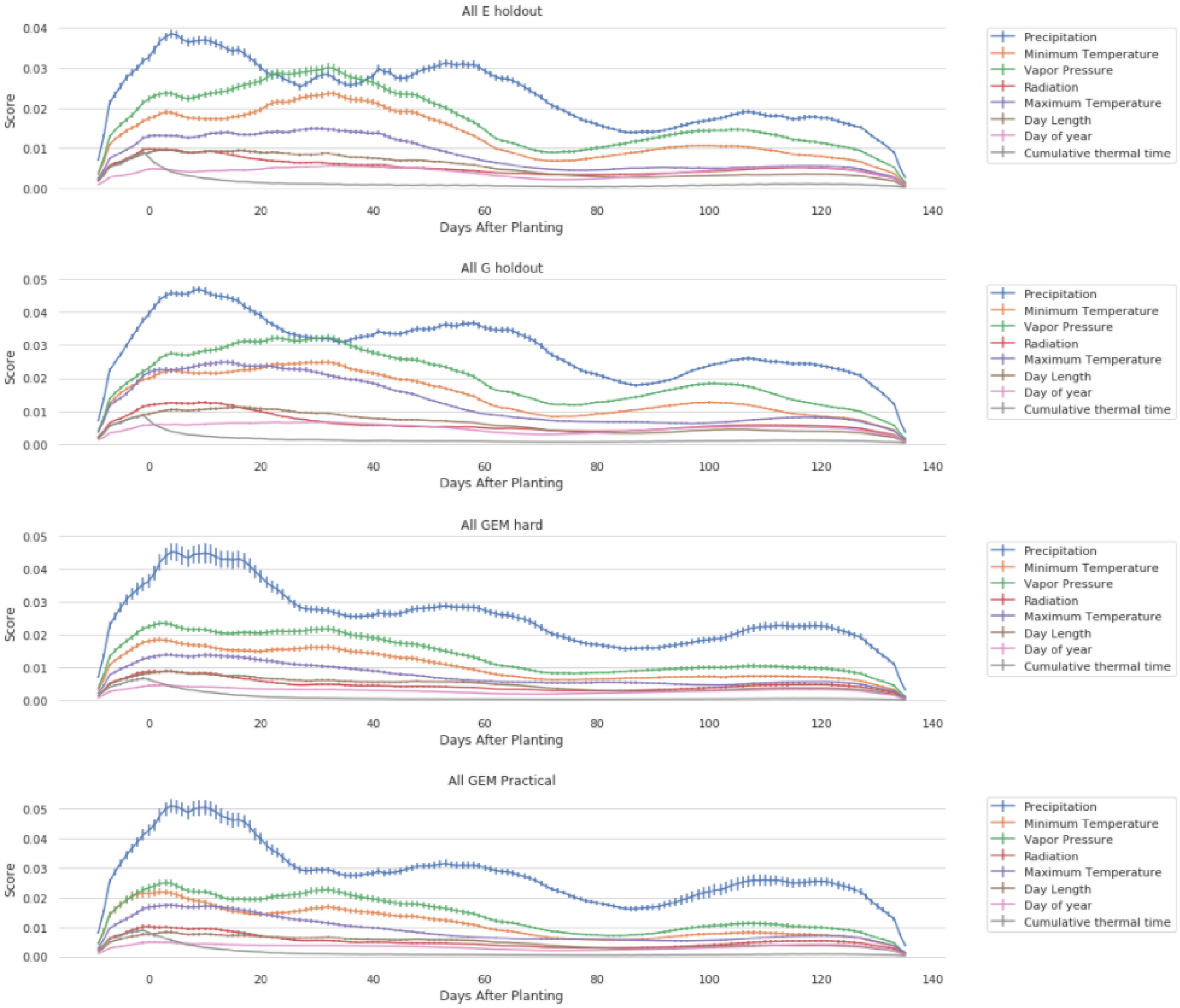


Supplemental Figure S6. Daily Saliancy map weather scores. Values are the average across all splits with historical pre-training included. Error bars are the standard error across splits.

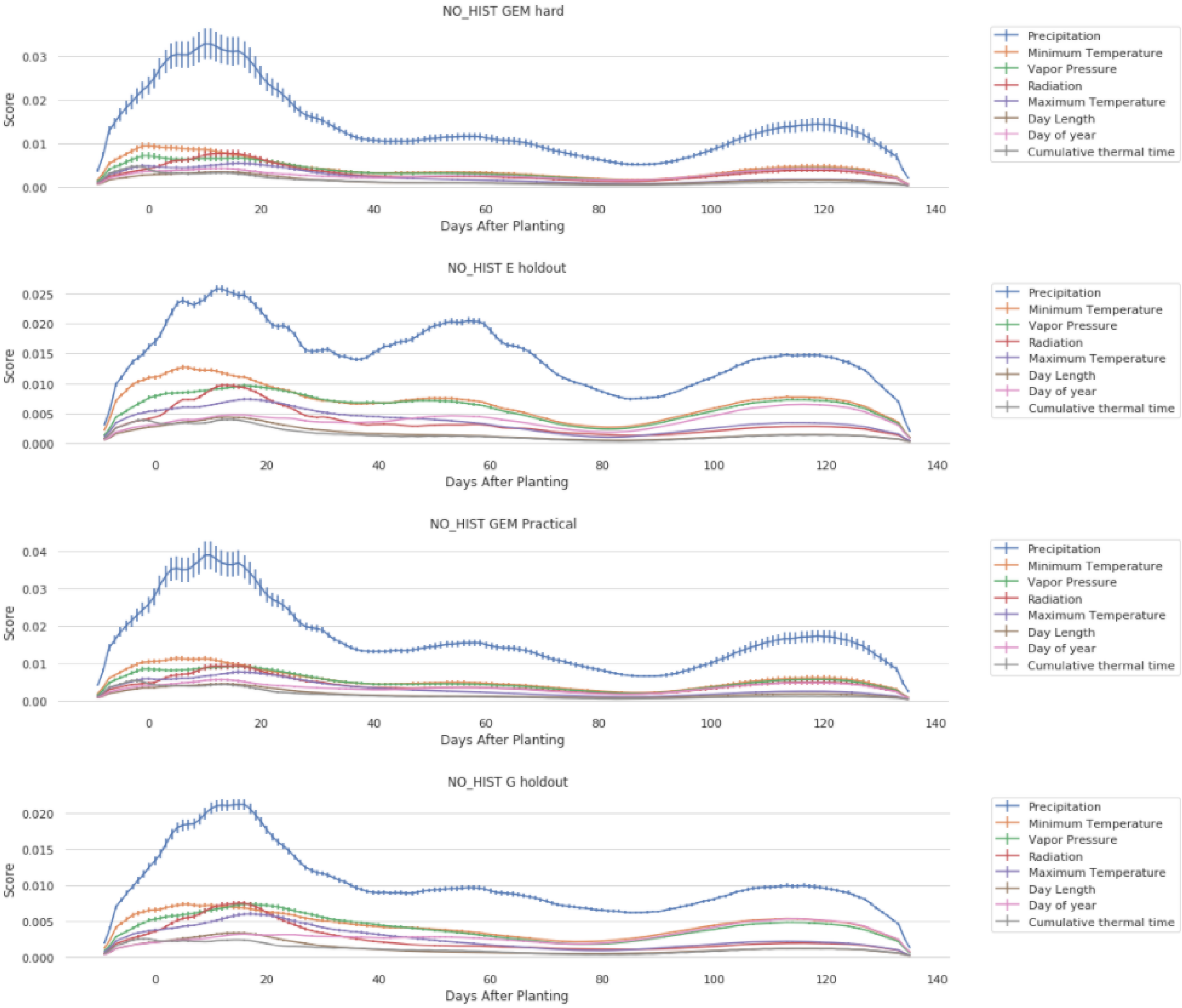


Supplemental Figure S7. Daily Saliancy map weather scores. Values are the average across all splits without historical pre-training included. Error bars are the standard error across splits.

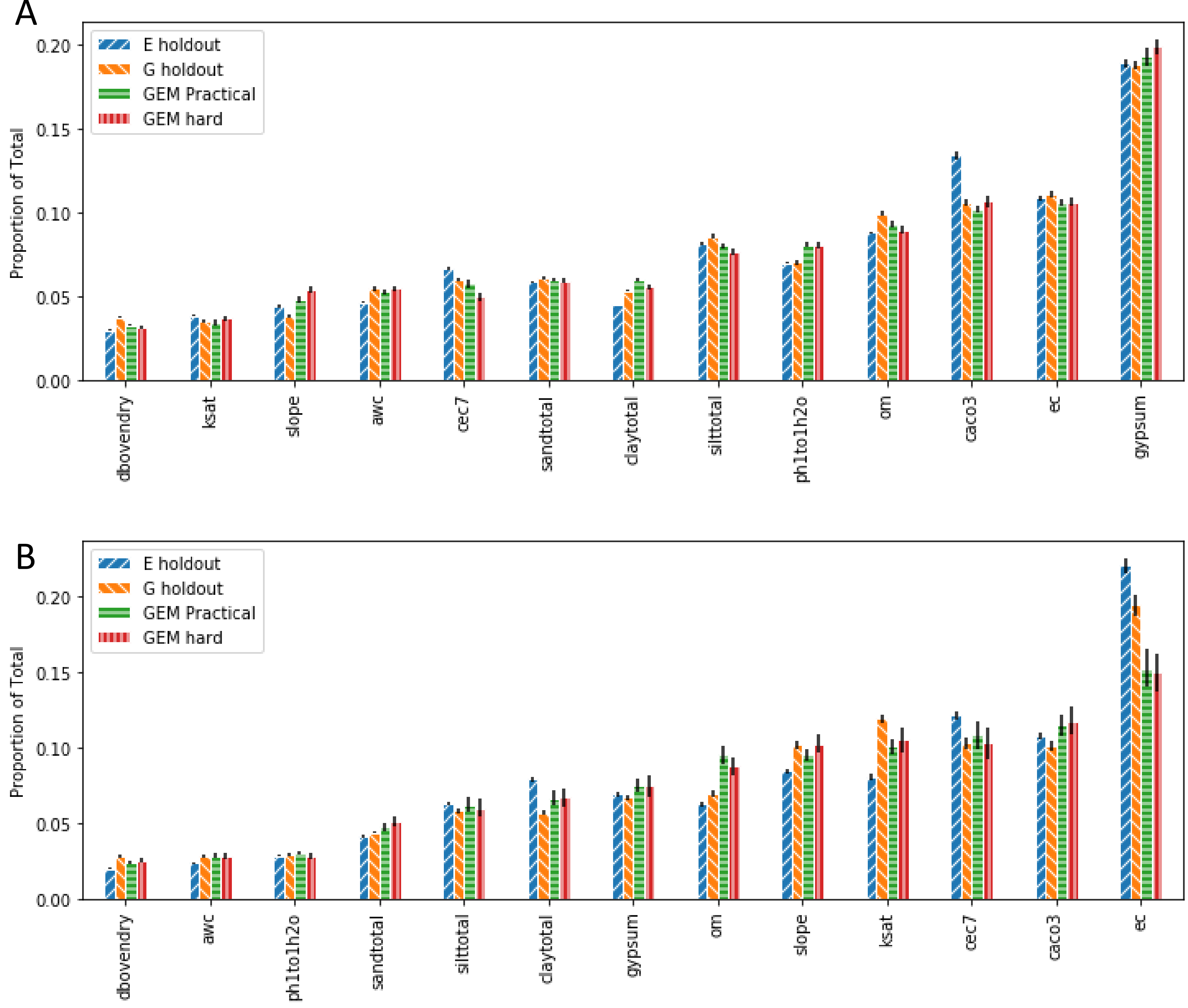


Supplemental Figure S8. Proportion of soil-related saliency map scores attributable to different factors. A) After pre-training on historical data, B) No pre-training on historical data. Factor names: dboverndry = oven dry soil weight, ph1to1h2o = pH of 1:1 soil-water mixture, awc = available water capacity, sandtotal = percentage of sand in soil, silttotal= percentage of silt in soil, claytotal = percentage of clay in soil, om = percentage of decomposed residue in soil, slope= soil slope, cec7= cation exchange capacity, caco3 = quantity of carbonate in soil, ec = soil electrical conductivity.

A

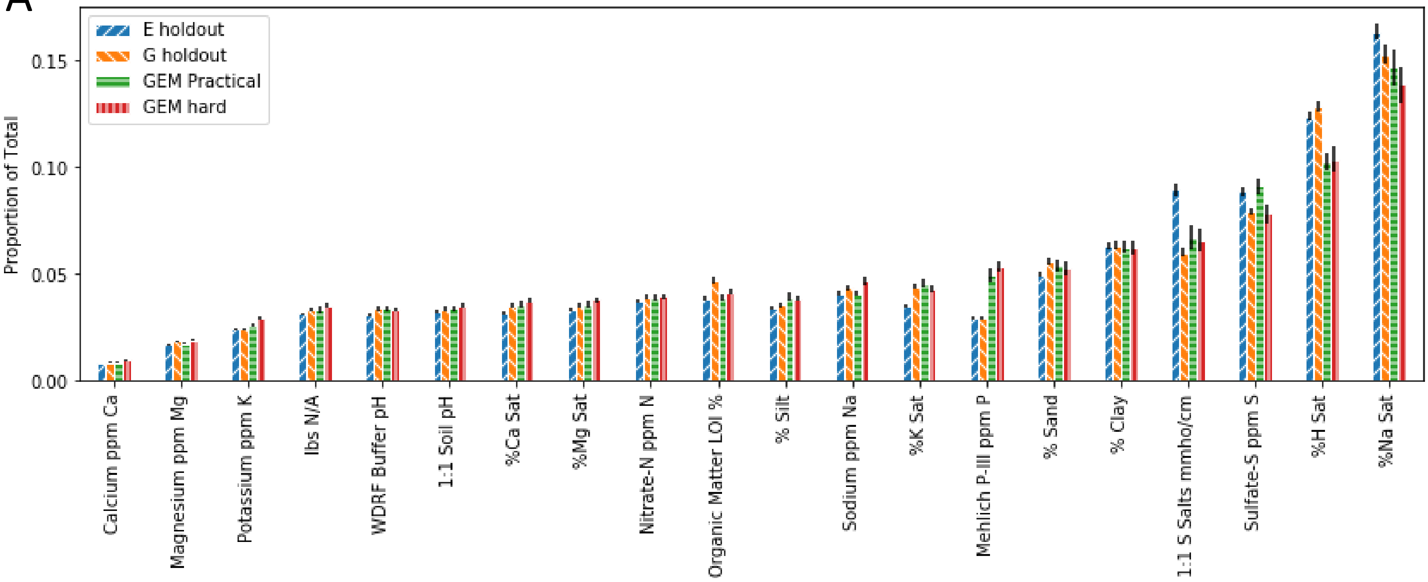


B

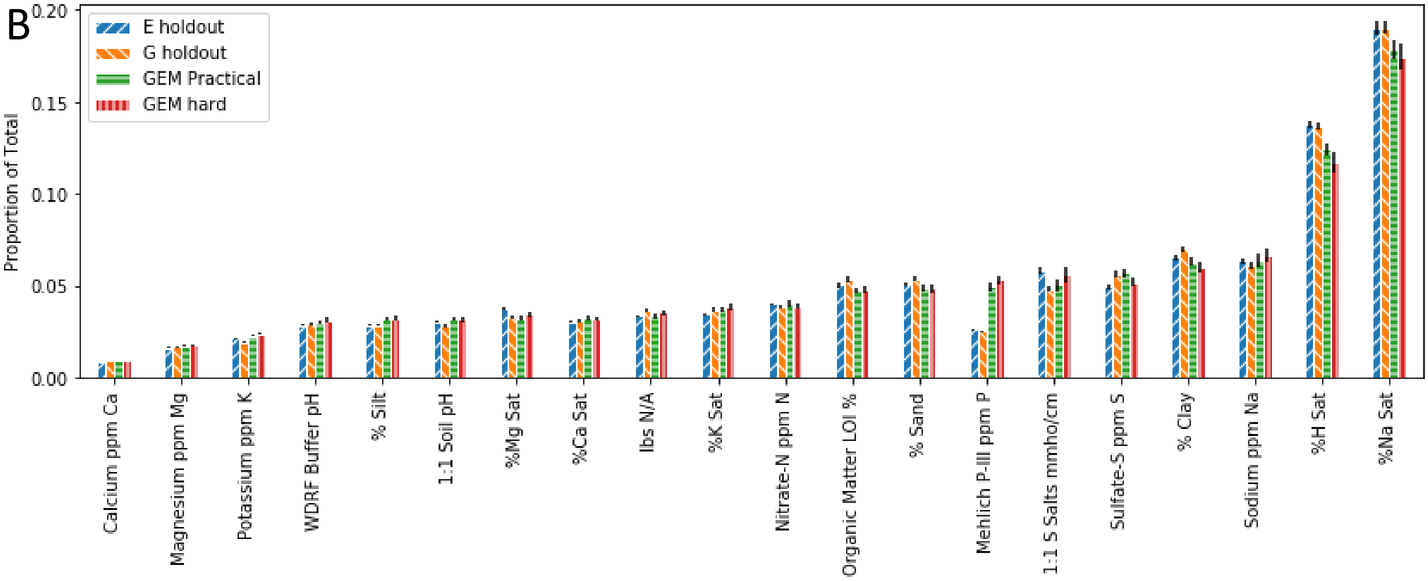


Supplemental Figure S9. Proportion of Fertility-related saliency map scores attributable to different factors. A) After pre-training on historical data, B) No pre-training on historical data.

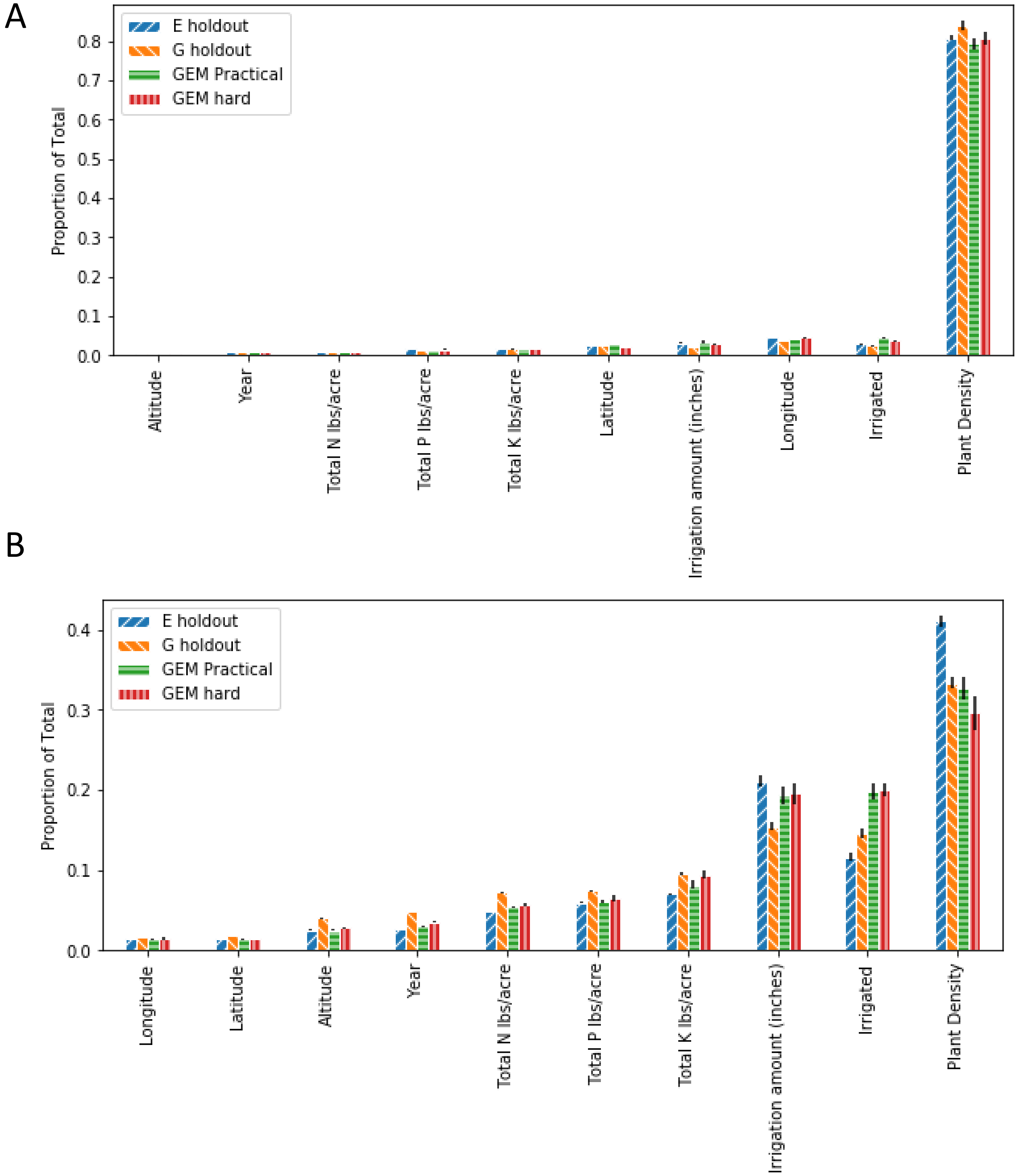


Supplemental Figure S8. Proportion of soil-related saliency map scores attributable to different factors. A) After pre-training on historical data, B) No pre-training on historical data.

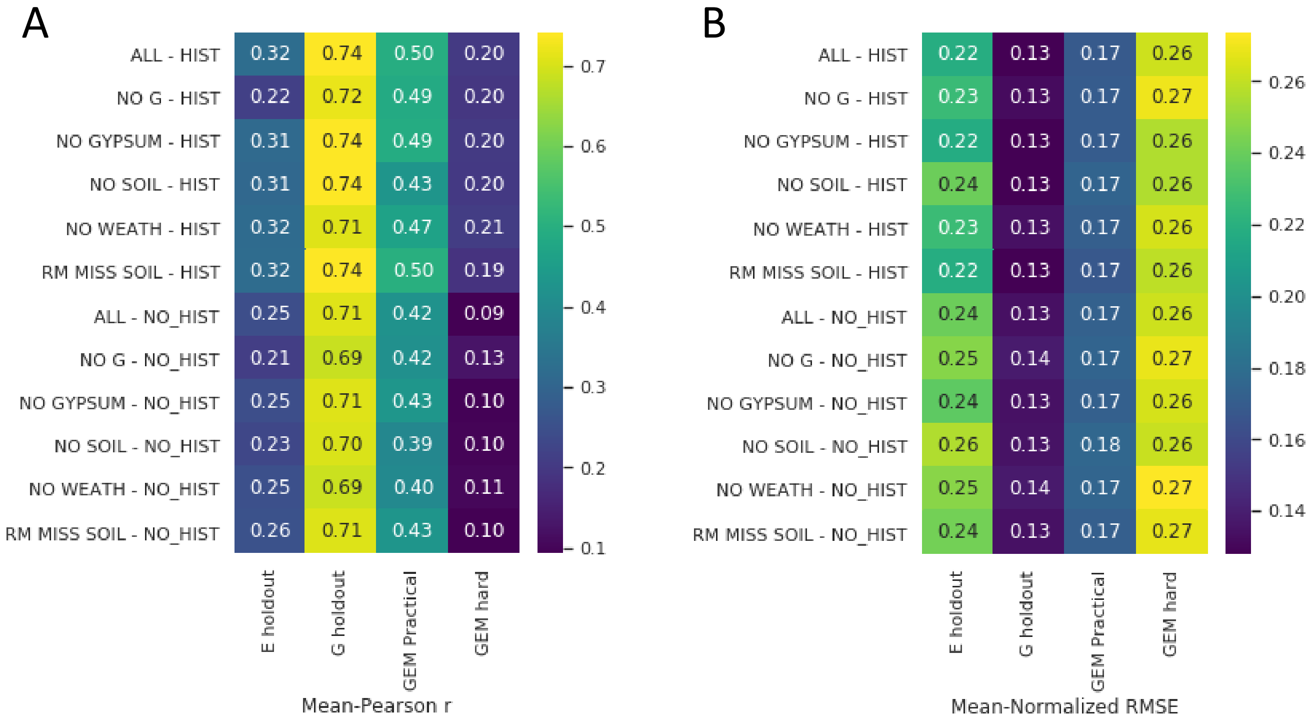


Supplemental Figure S9. Pearson r values (A) and normalized Root Mean Squared Error (RMSE) values (B) based on running the CNN model with different factors excluded. Note that excluding different factors from the model results in changes to the model's architecture, which make these comparisons potentially untrustworthy.
